# The transition from vision to language: distinct patterns of functional connectivity for sub-regions of the visual word form area

**DOI:** 10.1101/2023.04.18.537397

**Authors:** Maya Yablonski, Iliana I. Karipidis, Emily Kubota, Jason D. Yeatman

## Abstract

Reading entails transforming visual symbols to sound and meaning. This process depends on specialized circuitry in the visual cortex, the Visual Word Form Area (VWFA). Recent findings suggest that this word-selective cortex comprises at least two distinct subregions: the more posterior VWFA-1 is sensitive to visual features, while the more anterior VWFA-2 processes higher level language information. Here, we explore whether these two subregions exhibit different patterns of functional connectivity, and whether these patterns have relevance for reading development. We address these questions using two complementary datasets: Using the Natural Scenes Datasets (NSD; Allen et al, 2022) we identify word-selective responses in high-quality 7T individual adult data (N=8; 6 females), and investigate functional connectivity patterns of VWFA-1 and VWFA-2 at the individual level. We then turn to the Healthy Brain Network (HBN; Alexander et al., 2017) database to assess whether these patterns a) replicate in a large developmental sample (N=224; 98 females, age 5-21y), and b) are related to reading development. In both datasets, we find that VWFA-1 is more strongly correlated with bilateral visual regions including ventral occipitotemporal cortex and posterior parietal cortex. In contrast, VWFA-2 is more strongly correlated with language regions in the frontal and lateral parietal lobes, particularly bilateral inferior frontal gyrus (IFG). Critically, these patterns do not generalize to adjacent face-selective regions, suggesting a unique relationship between VWFA-2 and the frontal language network. While connectivity patterns increased with age, no correlations were observed between functional connectivity and reading ability. Together, our findings support the distinction between subregions of the VWFA, and portray the functional connectivity patterns of the reading circuitry as an intrinsic stable property of the brain.

## Introduction

Over two decades of cognitive neuroscience research have repeatedly located a region in the left ventral occipitotemporal cortex (VOTC) that responds selectively to text (for review, see Caffarra et al., 2021). This region, termed “the visual word form area” (VWFA), is consistently activated when words are presented visually. However, there are mixed findings regarding the specific functional properties of this region and the degree of selectivity it shows to language content compared with low level visual features (Dehaene and Cohen, 2011; Vogel et al., 2014; Caffarra et al., 2021; Yeatman and White, 2021). Recent evidence suggests that these inconsistencies may arise, at least in part, as a result of the VWFA not being a single uniform region but rather comprises a series of sub-regions, that differ in their response profiles and the level of computation they perform on the textual input (Lerma-Usabiaga et al., 2018; White et al., 2019; Yeatman and White, 2021). Specifically, it is suggested that the more posterior patch, VWFA-1, is sensitive to visual letter-shapes and orthographic information, and is often observed bilaterally. The more anterior VWFA-2 is sensitive to the language content of the visual text and is strongly left lateralized. These subregions differ from each other not only in their functional properties, but also structurally (Caffarra et al., 2021). First, these functional regions co-occur with different cytoarchitectonic areas in the visual cortex (Weiner et al., 2017a; Kubota et al., 2022). Second, diffusion MRI (dMRI) studies have shown they have different profiles of structural connectivity: VWFA-1 is connected via the vertical occipital fasciculus (VOF) to parietal regions involved in visual spatial attention, namely the inferior parietal sulcus (IPS). In contrast, VWFA-2 overlaps with cortical terminations of the posterior and long segments of the arcuate fasciculus (AF), which carry information between the temporal lobe, parietal lobe and the frontal lobe (Lerma-Usabiaga et al., 2018; Weiner and Yeatman, 2020; Kubota et al., 2022). Together, the emerging evidence suggests that, despite their proximity, these subregions are specialized for different components of the reading process. Here, we ask whether these subregions are also functionally associated with distinct *functional* networks in the brain.

### Functional connectivity of the VWFA

Resting state functional connectivity (rs-FC) is a methodology where the spontaneous fluctuations in the hemodynamic signal are compared between different regions in the brain. Distant regions that are highly correlated in time are considered to belong to the same network and share a common function (Biswal et al., 1995). Several studies have investigated the functional connectivity of the VWFA and found that it is highly correlated with frontal language regions (Koyama et al., 2010, 2011; Li et al., 2017, 2020), dorsal attention regions (Vogel et al., 2012) or both (Wang et al., 2014; Zhou et al., 2015; Chen et al., 2019). Importantly, these studies have typically looked at the VWFA as a single region, defined on a template using predefined coordinates or atlases (Koyama et al., 2010, 2011; Vogel et al., 2012; Wang et al., 2014; Schurz et al., 2015; Zhou et al., 2015; Li et al., 2017, 2020; Chen et al., 2019;

López-Barroso et al., 2020). It therefore remains unknown whether these connectivity patterns follow a similar division into subregions, and if so, is this unique to the reading network or a general principle of high-level visual cortex.

To tackle these questions, we capitalize on two complementary datasets. We first use the Natural Scenes Dataset (NSD) to define VWFA-1 and VWFA-2 at the individual level in high precision data, and investigate the functional connectivity patterns of each of these individually-defined subregions. Considering the evidence regarding the functional profiles of VWFA-1 and VWFA-2, we hypothesize that VWFA-1 will be strongly correlated with visual and attentional regions, primarily in the parietal lobe. In contrast, we expect VWFA-2 to be more strongly correlated with language regions in the left frontal lobe. We further define individual face-selective regions, to investigate whether this functional segregation is unique to reading related regions or applies to high-level visual processing more broadly.

We then turn to the Healthy Brain Network (HBN) dataset, a large pediatric sample consisting of hundreds of children and adolescents, to test whether a) these patterns replicate at scale and b) connectivity strength between the VWFAs and language regions is associated with reading development.

## Materials and Methods

### NSD dataset

The Natural Scenes Dataset (NSD; http://naturalscenesdataset.org) consists of 7T fMRI, anatomical MRI and diffusion MRI data for 8 healthy adult participants (6 females, 2 males) who viewed thousands of color images of natural scenes over multiple scanning sessions (see Allen et al., (2022) for a full description of the dataset). Additionally, each subject completed 20 sessions of resting state scans and six runs of a visual category localizer (“fLoc”) (Stigliani et al., 2015). NSD raw scans, minimally preprocessed data, and contrast maps for the functional localizer that were used for ROI definition, are all publicly available (https://cvnlab.slite.page/p/M3ZvPmfgU3/General-Information). See Table 1 for acquisition parameters.

**Table 1:** acquisition parameters for the two datasets, NSD and HBN. Note that HBN data were collected on multiple scanners.

|  | NSD | HBN |
| --- | --- | --- |
| Scanner | 7T Siemens <u>Magnetom</u> | 3T Siemens <u>TimTrio</u><br>3T Siemens Prisma |
| TR (ms) | 1600 | 800 |
| TE (ms) | 22 | 30 |
| Spatial resolution (mm) | 1.8 isotropic | 2.4 isotropic |
| Multiband acceleration factor | 3 | 6 |
| Run duration | 300s | 300s |
| Frames per run | 226 | 375 |
| Runs per subject | 20 | 2 |
| Total scan time | 100min | 10min |

### Data preprocessing and denoising

Data were minimally pre-processed using temporal resampling to correct for slice time differences, and spatial resampling to correct for head motion within and across scan sessions, EPI distortion and gradient non-linearities (for more details see Allen et al., (2022)). We then transformed each functional run to the individual cortical surface using the utility *nsd_mapdata* (https://github.com/cvnlab/nsdcode) and carried out all subsequent analyses on the native surface of each subject. Data denoising and correlation analysis were carried out using Nilearn v0.9.1 (Abraham et al., 2014). Data were denoised using a general linear model (GLM) with the following confound regressors and their temporal first derivatives, following the procedure described in (Yeo et al., 2011): six motion parameters, the white matter signal (averaged within a white matter mask eroded with 4 iterations), the ventricular signal, and the global signal averaged within a whole brain mask. We used an eroded white matter mask to minimize partial voluming between white and gray matter and reduce correlations between the white matter signal and the global signal (Power et al., 2017; Parkes et al., 2018). The global signal was included due to evidence showing it minimizes the dependency between connectivity measures and motion (Ciric et al., 2017). Although global signal regression can potentially increase negative correlations in the data (Saad et al., 2012), it has been shown to improve the specificity of positive correlations and reduce the influence of motion and respiratory artifacts (Murphy and Fox, 2017). Bandpass filtering was applied simultaneously to the signal and the confound regressors using a fifth-order Butterworth filter (0.008 - 0.1Hz). Lasly, we handled volumes with excessive motion by first flagging timepoints where the framewise displacement (FD) exceeded 0.25mm. We then created a binary (“spike”) regressor for each flagged timepoint. Runs where more than 10% of timepoints were flagged for motion, or where the mean FD exceeded 0.25mm, were discarded from analyses. This process resulted in excluding 4 runs for subj02, 3 runs for subj03, and 9 runs for subj05.

### ROI definition

We defined text-selective ROIs based on fLoc, a visual category localizer (Stigliani et al., 2015), where subjects viewed grayscale images of stimuli of different categories (e.g., text, faces, limbs, houses). Text-selective regions were identified on individual surfaces by comparing activations to text (pronounceable pseudowords) with activations to all other stimuli, except for numbers, thresholded at t>3. These contrast maps are included in the NSD data release. These contrast maps usually reveal multiple patches with several distinct activation peaks. To identify VWFA-1 and VWFA-2 we followed the guidelines in (White et al., 2023), focusing on patches within and around the occipitotemporal sulcus (OTS). VWFA-1 was defined as the posterior text-selective patch just anterior and lateral to area hV4, as suggested by Yeatman and colleagues (Yeatman et al., 2013). Manually labeled retinotopic ROIs were provided in the NSD release. VWFA-2 was defined as the next anterior, word-selective patch (using the same threshold). Activation clusters that overlapped the mid fusiform sulcus were not included. We next used the same contrast map (text > all stimuli except numbers) to define text-selective activations in the frontal lobe. These activations were generally found within the inferior frontal cortex (IFC), specifically centered around the inferior frontal sulcus (IFS) and inferior portion of the precentral sulcus (PCS), and are referred to as IFC-text hereafter. Lastly, for our control analyses using face-selective ROIs, we used the contrast map of faces vs. all other categories (t>3) to define face-selective regions along the fusiform gyrus, namely FFA-1 and FFA-2 (Chen et al., 2023). See Figure 1A for the ROI delineation process.

**Figure 1.**
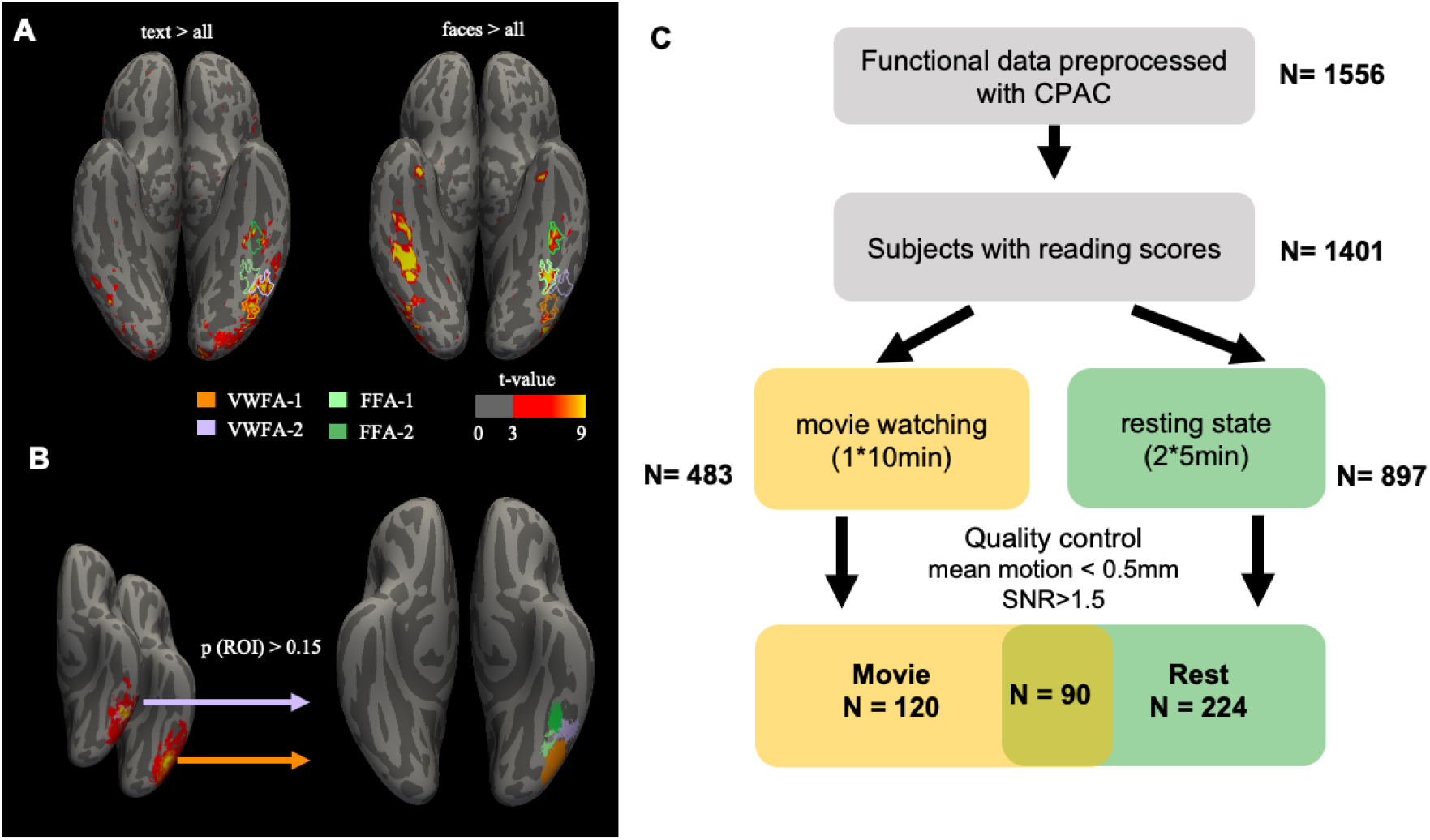
(A-B) Definition of category-selective ROIs. (A) For NSD subjects, VWFA-1 and VWFA-2 were defined using thresholded maps of text-selective responses (text > all other categories; t>3) on the individual cortical surface. Similarly, FFA-1 and FFA-2 were defined using the contrast faces > all other categories, with the same threshold. (B) Definition of group ROIs used for analysis of HBN data. Probabilistic maps were created by transforming individual ROIs defined in an independent sample of 28 adults to the fsaverage template. These maps were thresholded at p>0.15. The same procedure was used to define both text-selective and face-selective ROIs (C) Schematic of the sample selection process from the HBN database.

### Seed-based correlation analysis

For each seed ROI we extracted the mean time series, averaged across all vertices comprising the given seed ROI. We then calculated Pearson’s correlation coefficient between the seed timeseries and the timeseries of all other vertices on the cortical surface to generate a correlation map for each hemisphere. R values were converted to z-values using the Fisher transform to account for the non-normal distribution of correlation coefficients. We repeated this process separately for each run and then averaged the resulting correlation maps across runs, for each seed ROI for each subject. We used paired samples t-test to examine the difference between the correlation maps of VWFA-1 and VWFA-2. Next, to examine these correlations at the group level, we transformed each subject’s native surface to the fsaverage template space (using *nsd_mapdata* utility) and created group correlation maps across subjects (N=8). This whole-brain analysis provided us with a qualitative understanding of the spatial distribution of correlation patterns of VWFA-1 and VWFA-2 throughout the brain. All analyses were carried out using Nilearn v0.9.1 and visualized with Freeview v7.3.2 (https://surfer.nmr.mgh.harvard.edu/).

### ROI-to-ROI correlation analysis

To directly test the hypothesis that VWFA-2 is more strongly connected to language regions in the frontal lobe, we ran an ROI-to-ROI analysis where we quantified the connectivity strength between pairs of ROIs. Specifically, we calculate Pearson’s correlation coefficient between the timeseries of VWFA-1 and IFC-text, and VWFA-2 and IFC-text.

### HBN dataset

The Healthy Brain Network (HBN) database consists of neuroimaging and phenotypic data from over 1,500 participants ranging from 5 to 21 years of age from the greater New York City area (Alexander et al., 2017). For the current project, we capitalized on the availability of preprocessed functional MRI data (N=1556; publicly available on s3://fcp-indi/data/Projects/HBN/CPAC_preprocessed_Derivatives/). See Table 1 for acquisition parameters.

### Participant selection

As expected in large pediatric samples, some scans were of low quality or incomplete. We employed a strict quality control threshold to focus our analyses on high-quality, low-motion data. Specifically, we only included scans where the mean FD was lower than 0.5mm in each of the functional runs, and the signal-to-noise-ratio (SNR) was higher than 1.5. Figure 1 describes the full selection process that led to a final sample of N=224 participants (98 females; 126 males) who completed two high-quality resting state runs in addition to reading evaluation. In addition, we analyzed a partially overlapping sample of N=120 participants (48 females; 72 males) who completed a single high quality movie-watching run (equivalent in duration to the two resting state runs) and reading assessments. Inspection of the distribution revealed that despite the strict selection criteria, the resulting sample spanned children of all ages (as young as 6 years old) and a wide range of reading ability. Of the 224 participants, 44 had a dyslexia diagnosis, 115 had an ADHD diagnosis, and 26 had both diagnoses.

### Preprocessing

Functional data preprocessed using the CPAC package (Craddock et al., 2013) were obtained from the publicly available fcp-indi bucket on Amazon Web Services (s3://fcp-indi/data/Projects/HBN/CPAC_preprocessed_Derivatives). Standard preprocessing steps began with deobliqing and reorienting the anatomical scan. The functional scans were then motion corrected, skull stripped and coregistered to the anatomical scan. Data were resamped to 2^*^2^*^2mm voxel resolution and warped to MNI standard space. Denoising included the following regressors: six motion regressors, their quadratic terms, delayed regressors and quadratic delayed regressors; polynomial detrending with linear and quadratic terms. aCompCor was used to remove the first 5 components of the white matter and CSF components. Bandpass filtering was applied to maintain the frequency range (0.01 - 0.1Hz). Timepoints with excessive motion (FD > 0.5mm) were flagged for spike regression, in addition to the previous timepoint and the following 2 timepoints. Lastly, data were demeaned and normalized such that all signal timeseries are represented as z-scores.

### ROI definition

As no category localizer was available for this dataset, we used template ROIs defined in standard Freesurfer fsaverage template space. ROIs were defined as probability maps on an independent sample of adults (N=28; see Kubota et al., (2022)), who completed the same category localizer used by the NSD project (Stigliani et al., 2015). VWFA ROIs were defined as text-selective activations of text>everything else but numbers, thresholded at t>3. Then individual ROIs were transformed to fsaverage space to generate a probabilistic map where each vertex equals 1 if there was a perfect overlap across subjects in that location, or zero if none of the subjects had that ROI in that location (same procedure as in Rosenke et al., (2021). These probabilistic maps were thresholded at 0.15 (see Figure 1). The same procedure was used to define the face-selective ROIs, FFA-1 and FFA-2. To avoid confusion with the individual ROIs used for NSD, we refer to these group ROIs as gVWFA-1, gVWFA-2, gFFA-1 and gFFA-2.

We used a more lenient threshold of t>2 to define the frontal text-selective ROI, as text-selective activations were weaker in the frontal cortex than the occipitotemporal ROIs, and less consistent in location across individuals (N=28, Kubota et al., 2022). Then the group ROI probability map was thresholded at 0.15, smoothed with 1 step and anatomically constrained to the sulcus, to generate the template gIFC-text ROI.

### Seed-based correlation analysis

As a first step, all data were transformed from volume to fsaverage surface space using nilearn vol2surf function. We then followed the same procedure described above for the NSD dataset, namely, using gVWFA-1, gVWFA-2 and gIFC-text as seeds for whole-brain correlation analysis. We generated correlation maps where each vertex has the value of the z-transformed Pearson’s correlation value with the seed ROI timeseries. Statistical analyses were carried out across subjects. Specifically, we used paired samples t-test to examine the difference between the correlation maps of gVWFA-1 and gVWFA-2 across subjects.

### ROI-to-ROI correlation analysis

We followed the same procedure described above for the NSD database, this time using the template ROIs. We calculated Pearson’s correlation coefficient between the timeseries of gVWFA-1 and gIFC-text, and gVWFA-2 and gIFC-text. These ROI-to-ROI correlation values were used for the statistical analyses described below.

### Examining connectivity patterns of face selective ROIs

To determine whether the effects we found were specific to the reading ROIs or reflect a general functional organization of the ventral cortex, we repeated all the correlation analyses reported above while using face-selective ROIs as seeds, namely gFFA-1 and gFFA-2. We then ran an ROI-to-ROI correlation analysis, calculating the correlations between the timeseries of gFFA-1 and gIFC-text, and gFFA-2 to gIFC-text. We used gIFC-text as a frontal ROI for this analysis since no face-selective activations were identified in the inferior frontal cortex. Lastly, we ran a linear mixed effects model using the lme4 package in R (Bates et al., 2015), to assess how functional connectivity strength between each ventral ROI and IFC-text is influenced by visual category (text vs. faces), location (posterior OTS vs. mid-OTS) and their interaction. We then added to the model age, gender and participant motion in the scanner (average FD) as additional fixed effects, to examine how these subject specific characteristics interact with functional connectivity strength. All models included random intercepts for each subject. We compared models using the anova function in R to determine which of the factors and interactions made a significant contribution to the model fit.

### Brain-behavior association

To investigate whether functional connectivity patterns are associated with reading development, We fit ordinary least squares regression models with either reading skill or age as the dependent variable and functional connectivity strength, gender and participant motion in the scanner (average FD) as predictors. Reading skill was measured using the single word reading subtest of the Wechsler Individual Achievement Test (WIAT). Unless otherwise noted, age standardized scores were used, to tease apart the effects of reading skill from age.

## Results

### VWFA-2 is strongly correlated with frontal language regions across individuals

We first evaluated seed-based functional connectivity maps in individual subjects in the NSD database. While VWFA-1 was strongly correlated with ventral and dorsal visual regions in the occipital, temporal, and parietal lobes, VWFA-2 showed stronger correlations with regions in the frontal lobe, including the precentral sulcus (PCS), inferior frontal sulcus (IFS) and inferior frontal gyrus (IFG), bilaterally. Fig 2A shows the connectivity maps for a single participant (see Supplementary Figures 1-3 for the entire sample). Strong functional connectivity between VWFA-2 and regions of inferior frontal cortex (IFC) were clear at the individual level, although the exact location and size varied. To determine locations where functional correlations are significantly stronger with VWFA-2 compared with VWFA-1, we next calculated a paired samples t-test comparing the correlation maps within each individual subject. (Figure 2C and supplementary Figures 4-6). These maps confirm that VWFA-2 shows significantly stronger correlations with the IFC at the individual level for most participants (6 out of 8). While the magnitude and exact location of this effect are variable across participants, unthresholded maps (supplementary Figures 4-6) reveal remarkable consistencies in the general spatial pattern of the VWFA-2/VWFA-1 comparison across the eight participants. Of note, one of the subjects where the effect was not observed had poor data quality due to substantial motion (9/20 runs had to be excluded from analyses due to high motion, see Methods for exclusion criteria).

**Figure 2:**
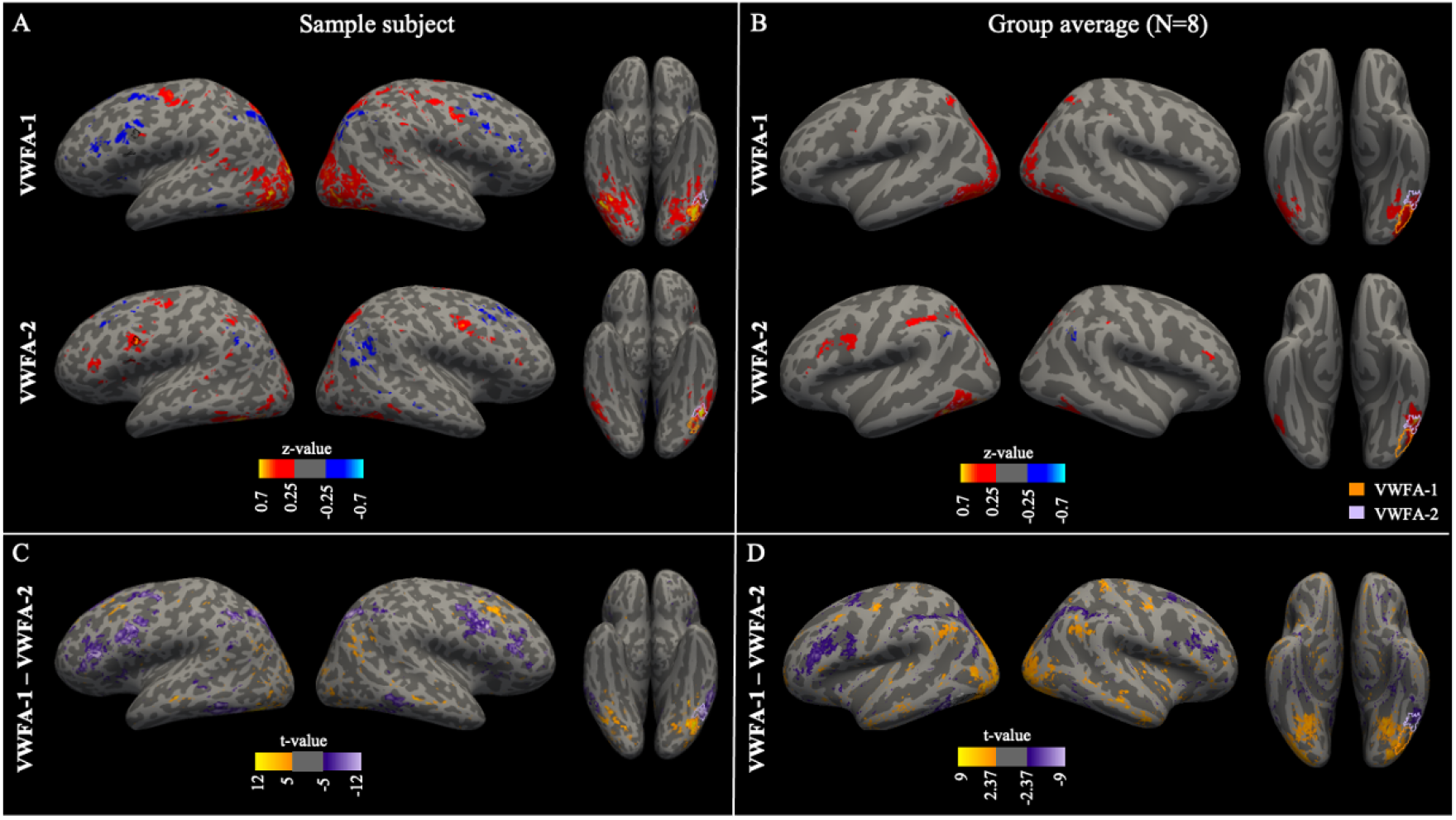
VWFA-1 and VWFA-2 show distinct patterns of functional connectivity at the individual level. (A-B) Seed based whole brain functional connectivity maps for VWFA-1 (top row) and VWFA-2 (bottom row) overlaid on inflated cortical surfaces, for a representative subject from the NSD dataset (subj03; Panel A) and averaged across the NSD sample (N=8; Panel B). Maps depict Fisher z-transformed Pearson correlation coefficients. (C-D) Significant differences in connectivity strength quantified using paired samples t-test for subj03 (Panel C) and averaged across the sample (D). In both C and D, positive values represent stronger connectivity with VWFA-1, negative values represent stronger connectivity with VWFA-2. Note the different threshold used for within subject analysis (t > 5, p < 0.0001, n=20 runs) and group analysis (t>2.365, p = 0.05 uncorrected, N=8 participants). In all panels, seed ROIs are shown on the ventral surface in orange (VWFA-1) and lilac (VWFA-2) contours. See supplementary Figures S1-3 for correlation maps of all individual NSD participants.

To summarize the individual effects observed in NSD data, we transformed the individual correlation maps into fsaverage template space and averaged the correlation maps across participants. This group analysis revealed a consistent patch spanning the posterior part of the IFG, PCS, and PCG, that showed stronger correlations with VWFA-2 compared with VWFA-1 (p

< 0.05, uncorrected). Thus, the high-precision, 7T NSD data revealed different patterns of functional connectivity between functionally defined VWFA-1 and VWFA-2 at the individual level, and template-based group average of these data indicated consistency in this effect across the eight participants.

After establishing that VWFA-2, compared with VWFA-1, is more strongly correlated with inferior frontal cortex in high-quality *adult* data in NSD, we turned to the HBN database to determine whether this pattern generalizes at scale in hundreds of individuals, and, importantly, whether it can be observed in a *developmental* population. To this end, we repeated the same seed-based analysis using template-based ROIs (see Methods) and compared the correlation maps of gVWFA-1 and gVWFA-2 across the participants in the HBN sample (Figure 3). These maps revealed, again, that gVWFA-1 is more strongly correlated with regions in the ventral occipital cortex, bilaterally, and inferior parietal regions, while gVWFA-2 is more strongly correlated with frontal regions and lateral parietal regions, primarily in the left hemisphere. These patterns were consistent for resting-state data (N=224) and movie watching data (N=120).

**Figure 3:**
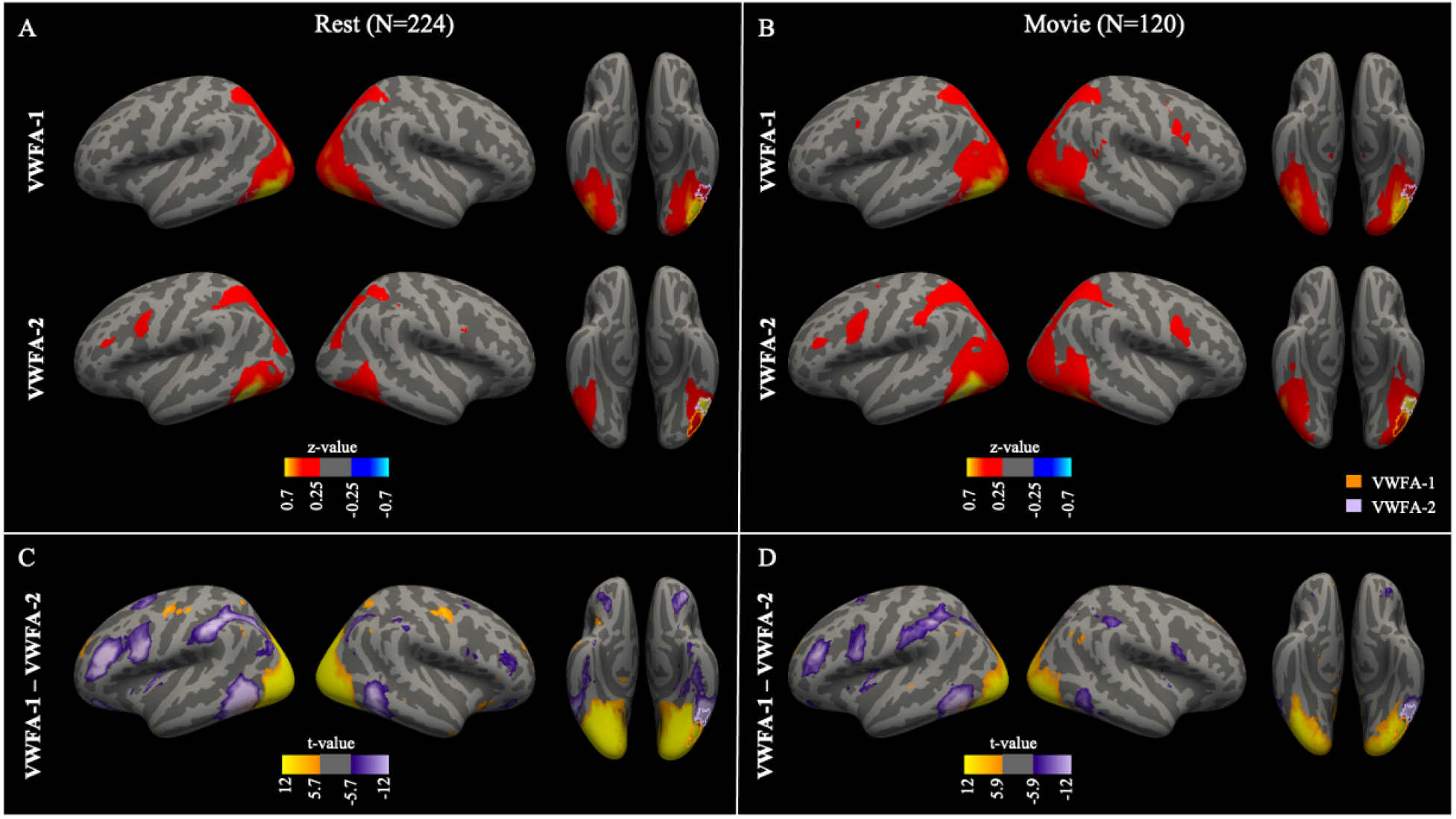
VWFA-1 and VWFA-2 show distinct patterns of functional connectivity in a large group of children and adolescents. (A-B) Seed-based whole brain functional connectivity maps for gVWFA-1 (top row) and gVWFA-2 (bottom row) overlaid on inflated cortical surfaces, for resting state (Panels A; N=224) and movie watching (Panel B; N=120). Maps depict Fisher z-transformed Pearson correlation coefficients. (C-D) Significant differences in connectivity strength quantified using paired samples t-test for resting state (Panel C) and movie watching (D). In both C and D, positive values represent stronger connectivity with gVWFA-1, negative values represent stronger connectivity with gVWFA-2. Maps are thresholded at p=0.01, Bonferroni corrected for the number of vertices (uncorrected p = 3e-8). In all panels, seed ROIs are shown on the ventral surface in orange (gVWFA-1) and lilac (gVWFA-2) contours.

### Text-selective regions in the frontal lobe show stronger functional connectivity with VWFA-2 compared with VWFA-1

We next set out to determine whether the correlations observed between VWFA-2 and the frontal lobe were specific to the language network, or rather represent an association with a more general dorsal attention network. To this end, we defined a text-selective region in IFC for each NSD participant, and used it as a seed for functional connectivity to inspect how the correlation pattern of that frontal region maps onto the ventral surface. This analysis revealed a striking correspondence between the connectivity map of IFC-text and the boundary between VWFA-1 and VWFA-2, both at the individual level and at the group level across the eight participants (Figure 4).

**Figure 4:**
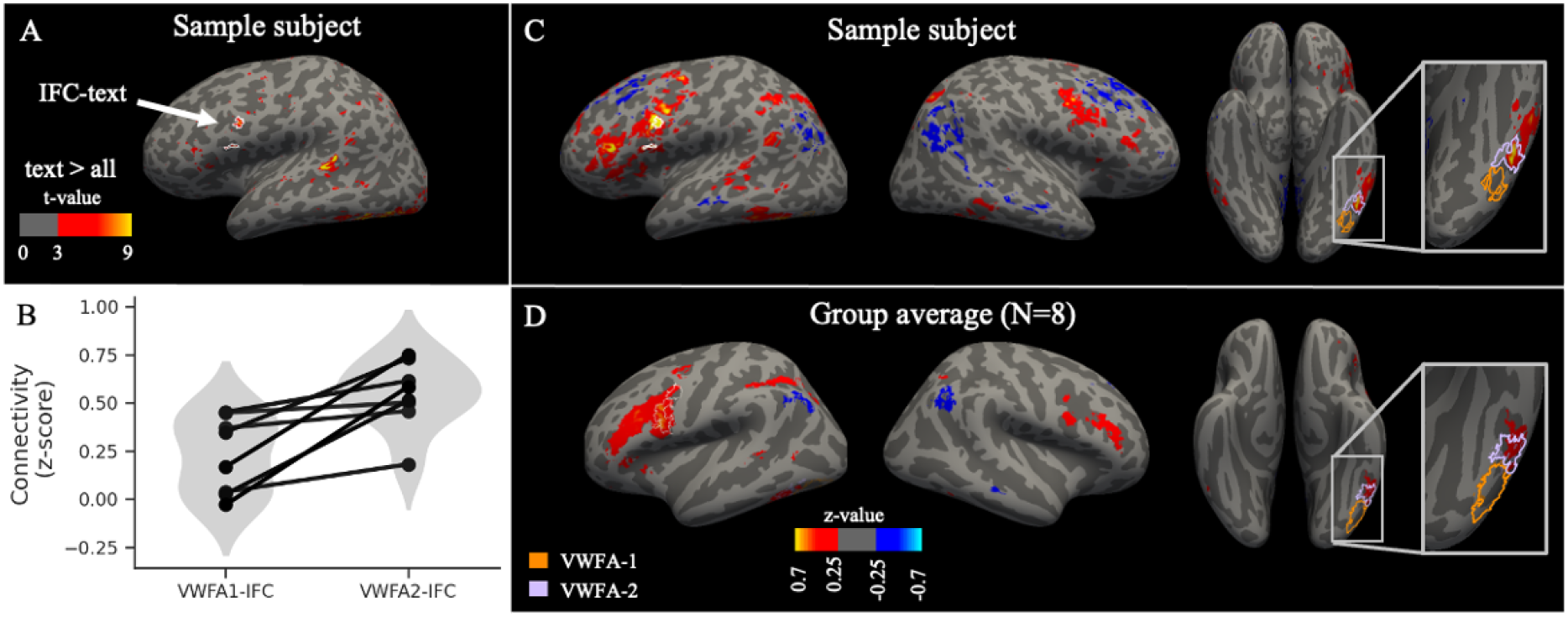
Frontal language regions show stronger connectivity with VWFA-2 compared with VWFA-1. (A) Text-selective activations (text vs. all other categories, t>3) for a single subject (subj03). The frontal activation patches selected as the frontal ROI (IFC-text) are delineated in white contour. (B) ROI-to-ROI correlation analyses revealed that the frontal ROI is more strongly connected with VWFA-2, compared to VWFA-1 (t(7) = 3.88, p = 0.0061). Each line represents a single participant (N=8). (C-D) Seed-based whole brain functional connectivity maps for the frontal text-selective region (IFC-text) overlaid on inflated cortical surfaces, for a representative subject (C; subj03) and averaged across the NSD sample (D; N=8). Maps depict Fisher z-transformed Pearson correlation coefficients. IFC-text for the participant in panel C is delineated by white contour. In panel D, ROI contours represent template ROIs.

To quantify this effect, we calculated the average correlation values between IFC-text and each of the two VWFA ROIs (Figure 4B). The resting state correlation between IFC-text and VWFA-2 was stronger compared with VWFA-1 (paired samples t(7) = 3.88, p = 0.0061). These specific correlations with the individually-defined IFC-text demonstrate that VWFA-2 is more strongly correlated than VWFA-1 with language-selective regions in the frontal lobe.

We then repeated the same analyses in the HBN dataset. Critically, as no individual localizer data is available for this dataset, we used template ROIs defined using text-selective activations in an independent dataset (see Methods). Even when using template ROIs as approximations for text-selective activations, our findings replicated at scale in this pediatric cohort (Figure 5): First, the correlation map of gIFC-text overlaps the boundary between gVWFA-1 and gVWFA-2, both for resting state data and movie watching data (Figure 5). Second, the ROI-to-ROI analysis confirmed that gIFC-text is more strongly correlated with gVWFA-2 compared with gVWFA-1, both when considering classical resting-state data(t(223) = 15.78, p < 1e-37) and movie watching data (t(119) = 12.28, p < 1e-22). This shows that the correlation between frontal language regions and VWFA-2 generalizes across populations (adults and children) and tasks (rest and movie watching).

**Figure 5:**
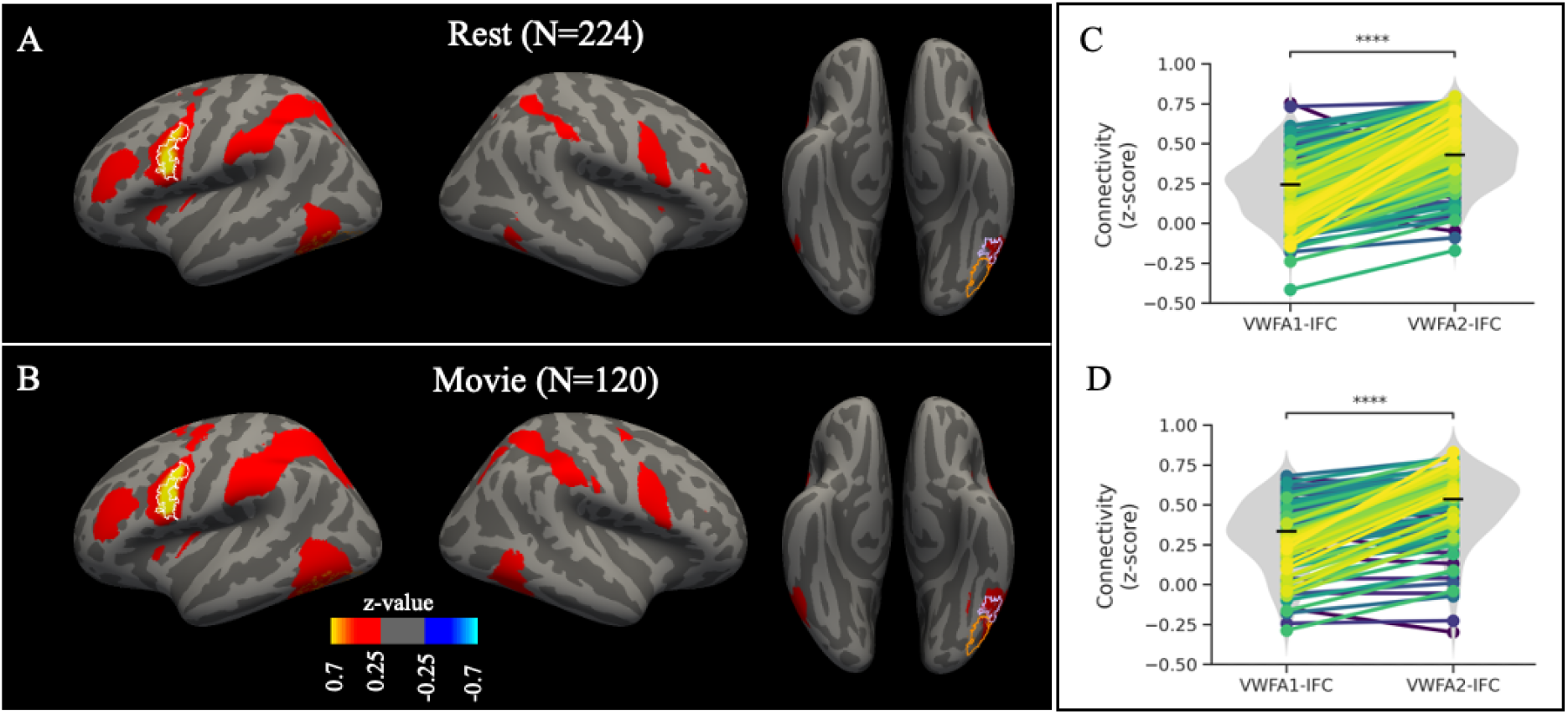
Frontal language regions show stronger connectivity with gVWFA-2 compared with gVWFA-1. (A-B) Seed-based whole brain functional connectivity maps for the frontal text-selective region (gIFC-text) overlaid on inflated cortical surfaces, averaged across the HBN sample for resting state data (A) and movie watching data (B). Maps depict Fisher z-transformed Pearson correlation coefficients. gIFC-text is shown in white contour. (C-D) ROI-to-ROI connectivity strength between gIFC-text and VWFA subregions. Connectivity is significantly stronger between gIFC-text and gVWFA-2 compared with gVWFA-1 (C: resting state, t(223) = 15.78, p < 1e-37; D: movie watching, t(119) = 12.28, p < 1e-22). Lines are color coded for the magnitude of the difference in connectivity strength between gIFC-text and gVWFA-1 compared with gVWFA-2, with larger differences in lighter colors. Black horizontal lines denote the median.

### Correlations between ventral occipital-temporal cortex and inferior frontal cortex are unique to the reading network

We next set out to determine whether these effects are indeed unique to the reading circuitry, or, alternatively, stem from a general organizational principle of ventral cortex. For example, we might hypothesize that the differences between VWFA-1 and VWFA-2 functional connectivity represent a general transition in connectivity between posterior and anterior regions of VTC. To this end, we first overlaid the face-selective ROIs (FFA-1, FFA-2) on the IFC-text connectivity map. We found that while the connectivity ‘hotspots’ overlap VWFA-2, they do not extend medially to the face-selective ROIs (Figure 6). To quantify this effect, we ran an ROI-to-ROI analysis on the resting state data (N=224), which revealed that FFA-2 does have stronger connectivity with IFC-text compared with FFA-1, but it is still significantly weaker than the effect observed for VWFA-2-IFS: The median connectivity between VWFA-2 and IFC-text was 0.43 (95%CI = [0.39, 0.45]), whereas the median connectivity between FFA-2 and IFC-text was 0.27 (95%CI = [0.23, 0.29]). A linear mixed effect model confirmed that functional connectivity values with the frontal lobe were significantly greater for text-selective regions compared with face-selective regions (*β*=0.025, t=2.35, p=0.0189) and for mid-OTS regions compared with posterior OTS regions (*β*= 0.049, t=4.59, p=5e-6). Importantly, the interaction between category and location was also highly significant (*β* = 0.135, t=8.89, p<2e-16), indicating that the correlation between VWFA-2 and the IFC was significantly stronger compared to the other ventral ROIs. These effects remained the same when participant age, gender and motion were included in the model as between subject variables. While age and subject motion each increased functional connectivity values overall (age: *β*=0.009, t=3.03,p=0.0027; motion: *β*=0.512, t=2.219, p=0.0275), gender was unrelated to functional connectivity strength (*β*=0.008, t=0.422, p=0.6736). To test whether any of these factors had a specific effect on VWFA2-IFC connectivity strength, we ran three separate models that included the three-way interaction between category, location, and each of the variables of interest (age, gender, and motion), and compared them to the baseline model that included only the main effects. None of the three-way interactions nor the two-way interactions with age, gender or motion were significant. An ANOVA comparing the models revealed that including these interactions did not improve the model fit beyond the contribution of the main effects of these variables. Running the same analysis on the movie watching data (N=120) revealed comparable effects (Category: *β*=0.044, t=3.05, p=0.0024; Location: *β*=0.033, t=2.27, p=0.0241) and a significant interaction showing that the correlation between IFC and VWFA-2 was significantly stronger compared with the other ventral ROIs (*β*=0.139, t=6,84, p=3e-11). Age, gender or motion did not make a significant contribution to the model in explaining functional connectivity values in the movie-watching data (all p > 0.05). In sum, this analysis showed that the IFC was more strongly correlated with VWFA-2 compared with the rest of the ventral ROIs, above and beyond the effect of location along the posterior-anterior axis. This effect is independent of subject age, gender and motion.

**Figure 6.**
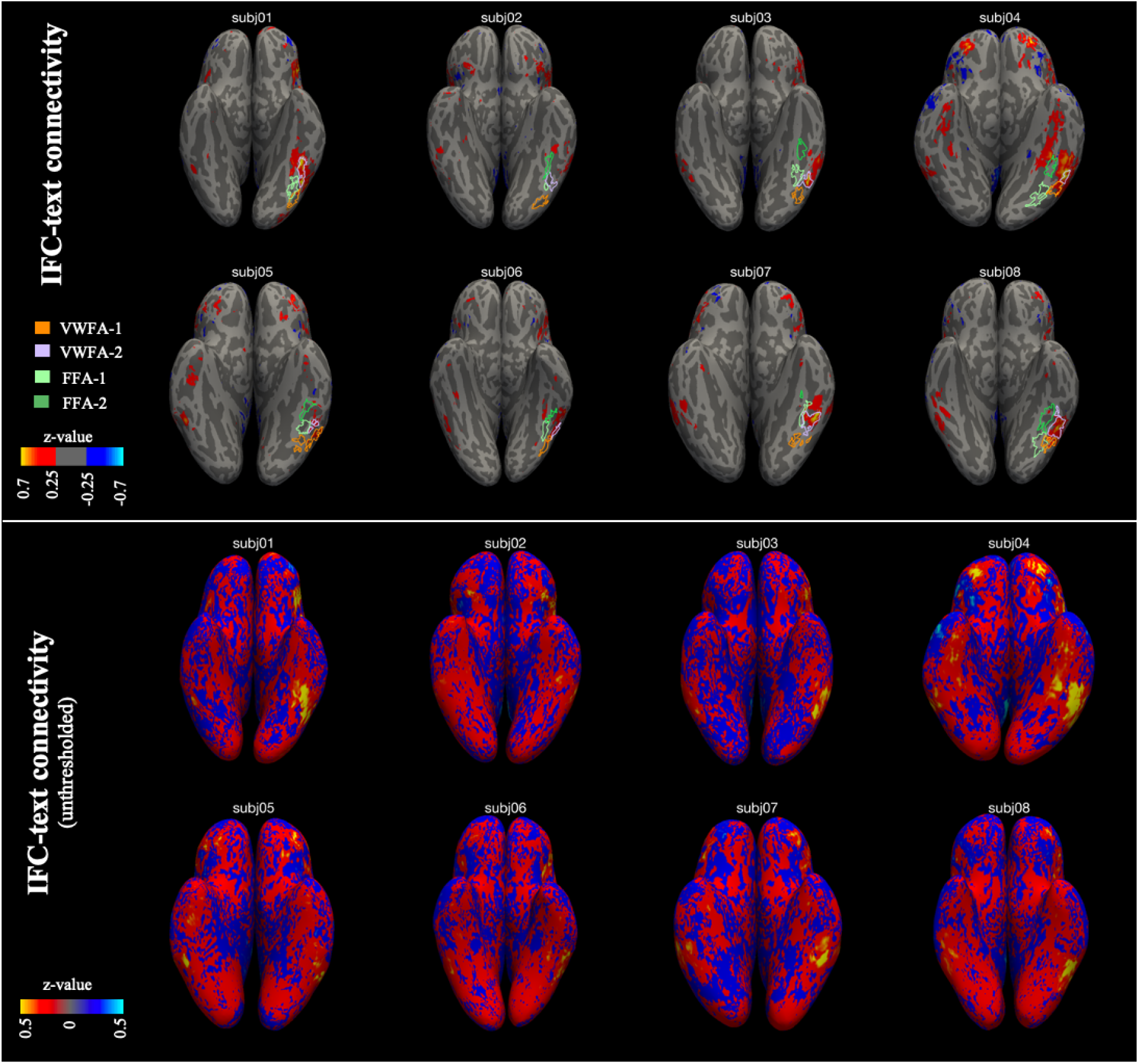
IFC-text is specifically correlated with VWFA-2. Shown are functional connectivity maps for the frontal ROI (IFC-text) on the ventral surface of individual subjects. Top panel-thresholded correlation maps with contours of individually defined category-selective ROIs. Bottom panel-unthresholded correlation maps.

To further explore the spatial distribution of the correlations of the face-selective compared to word-selective regions, we repeated our primary functional connectivity analysis using the face-selective ROIs as seeds. The resulting maps revealed that the correlations between FFA-1 and FFA-2 and IFC were significantly weaker than the correlation observed between VWFA2- and IFC-text (Figure 7 and Figure S7): using the same threshold, no patches of functional connectivity were observed in the frontal lobe. Lastly, we compared the whole brain correlation maps of VWFA-2 to those of FFA-2 to see how they differ spatially throughout the brain (Figure 7C-D). This revealed that VWFA-2 is more strongly correlated than FFA-2 with two large patches in the frontal lobe, as well as with a large area in the inferior parietal sulcus (IPS). In contrast, FFA-2 is more strongly correlated with regions in the superior temporal sulcus (STS) and the central sulcus.

**Figure 7:**
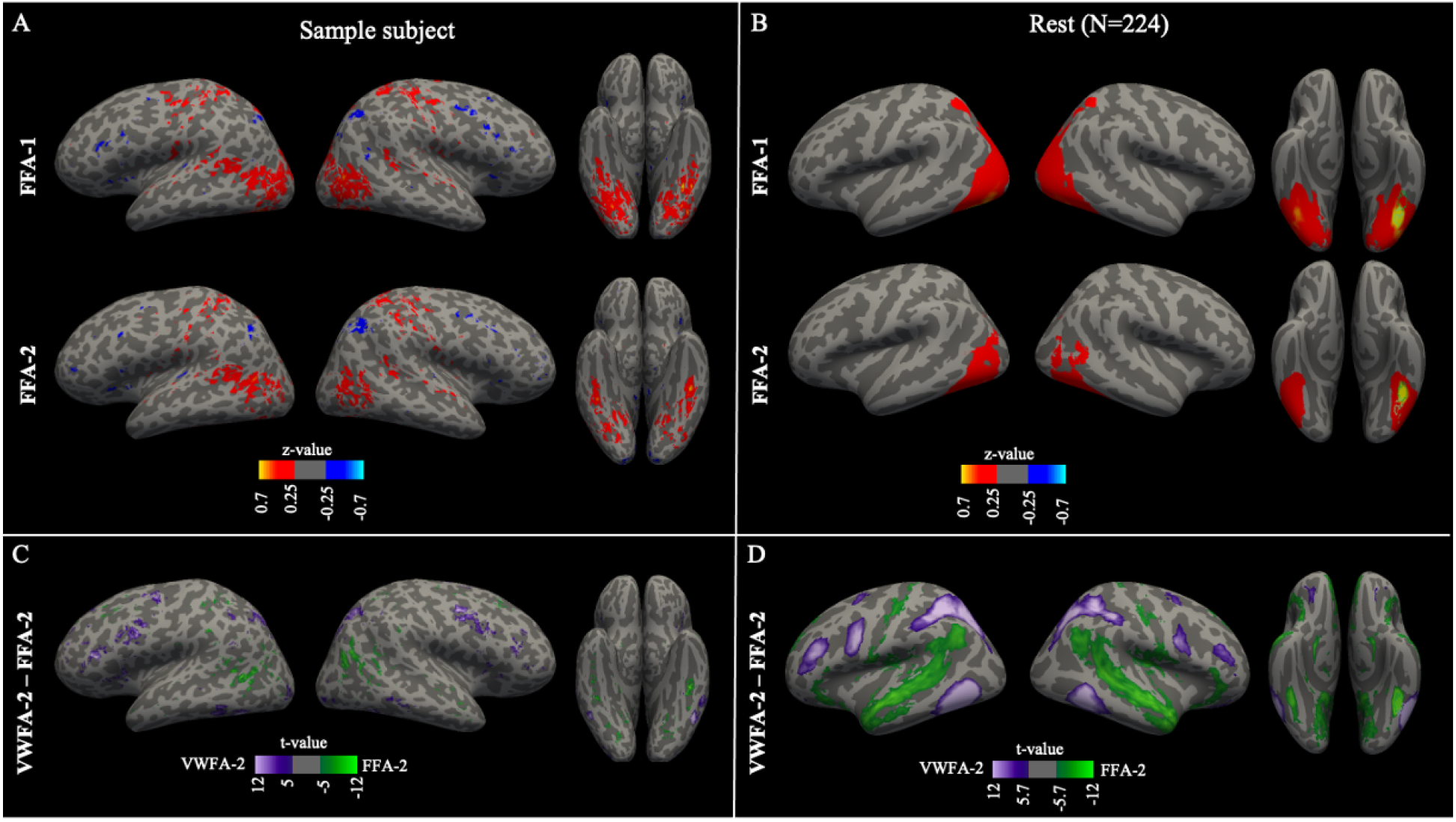
Face selective ROIs do not show the same correlations with the frontal lobe as the text-selective ROIs. (A-B) Seed-based whole brain functional connectivity maps for the face-selective ROIs, FFA-1 and FFA-2 overlaid on inflated cortical surfaces, for a single subject from the NSD dataset (subj03; A) and averaged across subjects from the HBN dataset (B). Maps depict Fisher z-transformed Pearson correlation coefficients. (C-D) T-test maps showing the difference in connectivity values between VWFA-2 and FFA-2, for the same single subject (C) and averaged across the HBN sample (D)

### VWFA functional connectivity develops with age, but is not related to reading ability

We next examined whether the functional connectivity strength between VWFA-1/2 and the frontal ROI is associated with reading development. We first used an ordinary least squares regression model, with age as the dependent variable, and functional connectivity values (VWFA1 to IFC-text and VWFA2 to IFC-text), gender, and average motion in the scanner (mean FD averaged across runs) as predictors. We found that functional connectivity strength between VWFA1 and IFC-text was a significant predictor of age (*β* = 4.569, p = 0.0009), such that the correlation between VWFA1 and the frontal language region increased with age. As expected, motion was highly predictive of age (*β* = -19, p= 0.00004), as younger children tend to move more in the scanner. In total, the model explained 11% of the variance in age (F = 7.1, p = 0.00002). We next created a second model predicting reading ability, which was operationalized as the age standardized single word reading subtest from the WIAT battery. We included the same predictors as in the previous analysis. The only significant predictor of reading skill was motion (*β* = -71, p= 0.0071), but the full model failed to reach significance and explained less than 5% of the variance in reading ability. In line with these results, the simple correlation between age and functional connectivity strength between IFC and VWFA-1 was significant (r=0.208, p = 0.002; Figure 8), while the other simple correlations between connectivity values and age or reading were non-significant (all p > 0.05). Applying the same regression models to the movie-watching data did not reveal any significant effects (all p> 0.1).

**Figure 8:**
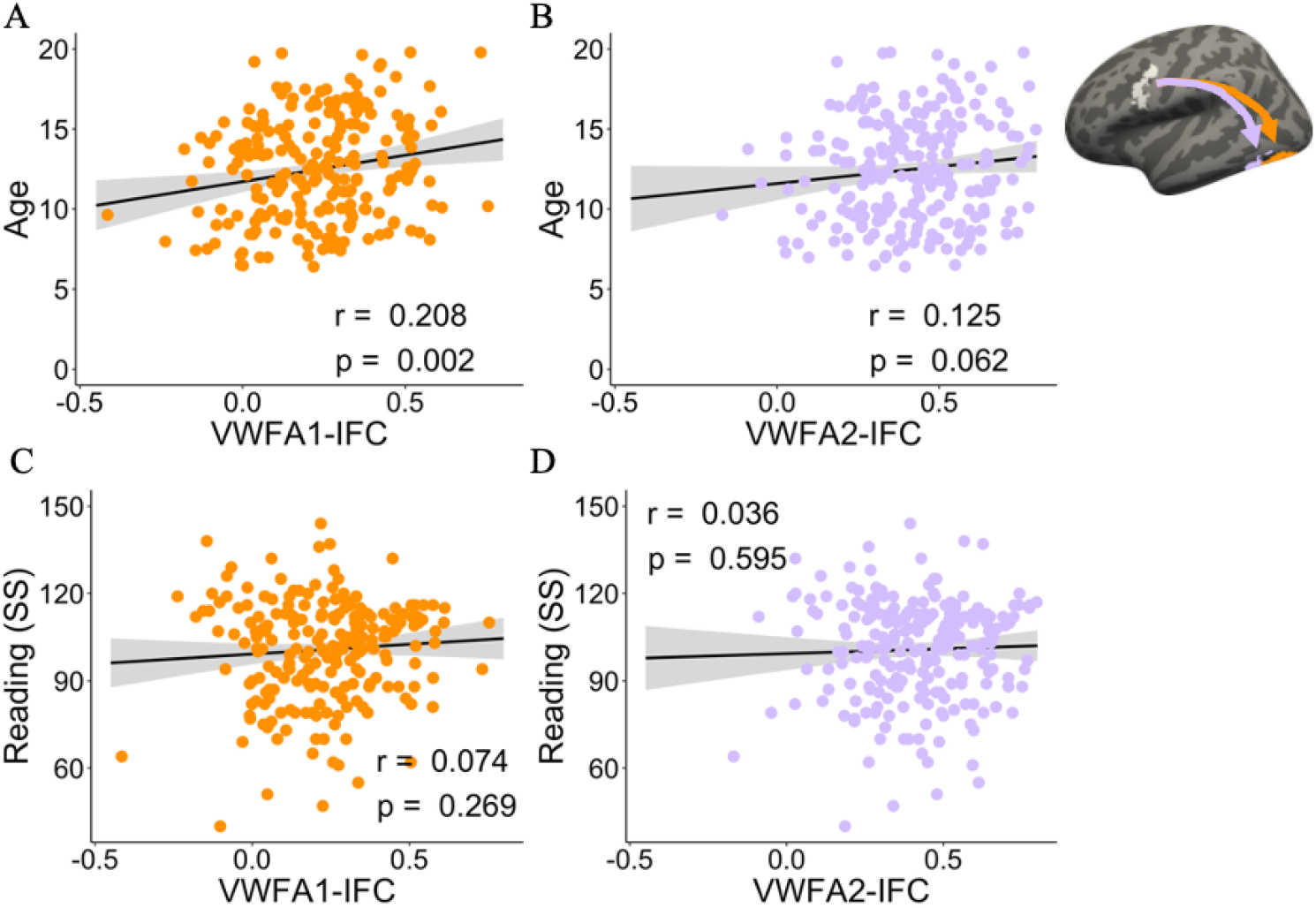
Functional connectivity strength is correlated with age, but not with reading ability. The ROIs are shown in the inset, with VWFA-1 in orange, VWFA-2 in lilac and the frontal language ROI (IFC-text) in white.

## Discussion

We found, in two independent datasets, that subregions of the VWFA show distinct patterns of functional coupling with the rest of the brain: The more posterior VWFA-1 is strongly correlated with visual and attentional regions in the occipital and parietal cortex, while the more anterior VWFA-2 is strongly correlated with language regions in the frontal lobe. We established this distinction first at the individual level in high quality adult data using individually-defined ROIs. We then confirmed that these patterns replicate at scale in a large sample of children and adolescents, and are independent of reading ability and task (movie watching compared with classical resting state). Importantly, we show that these patterns are unique to the reading circuitry and do not generalize to adjacent face-selective ROIs.

These findings are in line with the structural connectivity literature detailing the white matter connections of the VWFA (Yeatman et al., 2013; Bouhali et al., 2014; Lerma-Usabiaga et al., 2018). Recent studies reported that posterior and mid/anterior portions of the VTC are connected to different brain areas via separate long-range white matter pathways. Specifically, posterior regions were shown to be connected to the inferior parietal lobe through the VOF (Yeatman et al., 2014; Weiner et al., 2017b). In contrast, more anterior portions of the VTC were shown to connect with frontotemporal regions through the AF, a large tract spanning the temporal, parietal and frontal lobes, that has been repeatedly associated with reading and language abilities (Yeatman et al., 2011, 2012a, 2012b; Vandermosten et al., 2012; Saygin et al., 2013; Broce et al., 2015; Gullick and Booth, 2015; Skeide et al., 2016). Recent studies that combined a functional definition of the VWFAs, with diffusion MRI based fiber tractography, confirmed that functionally defined VWFA-1 and VWFA-2 overlap with endpoints of the VOF and the AF, respectively (Lerma-Usabiaga et al., 2018; Kubota et al., 2022). Our current findings corroborate this distinction, and reveal a correspondence between the functional and the structural connectivity of these adjacent subregions.

In light of these findings, we interpret our results as attesting to the different levels of analysis that are carried out in each subregion of the VWFA. Several studies have reported a hierarchy in the VTC, where sensitivity to lexical features and abstract language information increases along the posterior-anterior axis (Binder et al., 2006; Vinckier et al., 2007; van der Mark et al., 2009; Olulade et al., 2015; Taylor et al., 2019). Interestingly, a recent review found that studies that compare responses to text with responses to non-text stimuli, typically reveal text-selective activations in a wide range of locations along the posterior-anterior axis. In contrast, studies that localize the VWFA using lexical contrasts (e.g., comparing real words and pseudowords) consistently report a single, more anterior region (Caffarra et al., 2021). Indeed, when lexical contrasts were directly compared with orthographic contrasts in the same sample, orthographic contrasts revealed two patches, posterior and anterior, while lexical contrasts resulted in a single anterior patch (VWFA-2) (Lerma-Usabiaga et al., 2018). In addition, VWFA-1 and VWFA-2 were shown to have different response profiles when multiple words are presented in parallel, leading to the suggestion that VWFA-2 serves as a bottleneck for word processing (White et al., 2019). Taken together, the evidence suggests that VWFA-1 processes visual and orthographic information, which then undergoes further processing and integration with the language system in VWFA-2.

A crucial question is whether these distinct connectivity patterns are specific to the language network. An alternative explanation would be that this transition in connectivity patterns is an organizational principle of the VTC, that is independent of category-selectivity. For example, the motif of repeating, category-selective patches has been shown to be a general organizational principle of high-level visual cortex (Tsao et al., 2008; Grill-Spector and Weiner, 2014; Freiwald, 2020; Yeatman and White, 2021). In other words, it could be that there is a general posterior-anterior gradient of connectivity such that more anterior regions have stronger correlations with the frontal lobe. Our findings favor a view of the VWFA-2 as a distinct region with specific connectivity to language regions in the IFC: By examining the connectivity patterns of face-selective ROIs we confirm that a) VWFA-2 is strongly correlated with IFC-text above and beyond the effects of category-selectivity and location along the posterior-anterior axis. b) frontal text-selective regions within the IFC show a correlation peak that almost perfectly localizes VWFA-2, and does not extend medially to face-selective ROIs (Figure 6). We conclude that there is a unique relationship between VWFA-2 and language regions in the frontal lobe, further supporting its role in bridging visual-orthographic analysis and high-level language processing.

This view is in accordance with previous functional connectivity studies, which found that the VWFA is highly correlated with frontal language regions (Koyama et al., 2010, 2011; Li et al., 2017, 2020; Stevens et al., 2017). Other studies have reported that the VWFA is also correlated with dorsal attention regions in the parietal lobe (Wang et al., 2014; Zhou et al., 2015; Chen et al., 2019). These correlations with the dorsal attention network were taken to imply that the VWFA serves as a hub for visual attention that is not specific to reading (Vogel et al., 2012). Our approach of breaking down the VWFA into subregions resolves these discrepancies, by uncovering the different connectivity patterns of VWFA-1 and VWFA-2. Importantly, past studies have typically looked at the VWFA as a single region, defined on a template using predefined coordinates or atlas parcellations (Koyama et al., 2010, 2011; Vogel et al., 2012; Wang et al., 2014; Schurz et al., 2015; Zhou et al., 2015; Li et al., 2017, 2020; Chen et al., 2019; López-Barroso et al., 2020). Interestingly, when a single VWFA was defined in individuals using a functional localizer, higher connectivity values were found with language regions compared with a parallel analysis that used a group VWFA (Stevens et al., 2017). This highlights the importance of using individually defined ROIs when studying the fine tuned organization of text-selective cortical regions (Glezer and Riesenhuber, 2013). The current study takes a step further by delineating the distinct subregions of the VWFA in individuals and investigating their functional connectivity patterns separately.

Is functional connectivity of the VWFA related to reading ability? The evidence is mixed. As a group, struggling readers were shown to have decreased functional connectivity between the VWFA and frontal language regions (Schurz et al., 2015; Zhou et al., 2015; López-Barroso et al., 2020). Several studies also reported that FC strength is correlated with reading performance in children (Alcauter et al., 2017; Cross et al., 2021) and adults (Koyama et al., 2011; Achal et al., 2016; Li et al., 2017; Stevens et al., 2017; López-Barroso et al., 2020). However, these associations were found in inconsistent locations in the brain, for example, in the striatum (Achal et al., 2016), thalamus (Cross et al., 2021) and left IFG (Cross et al., 2021). Of the studies that focused on VWFA connectivity, some found associations between reading and FC strength between the VWFA and Wernicke’s area in the temporal lobe (Stevens et al., 2017), while others highlighted the connections between the VWFA and frontal language regions (Koyama et al., 2011; Li et al., 2017). Interestingly, these correlations with reading were observed in adults but did not generalize to children (Koyama et al., 2011; Li et al., 2017). In our current sample of children and adolescents (N=224) we did not find evidence for correlations between reading ability and functional connectivity of the VWFA. Interestingly, a recent large-scale study (N=313) found that while structural connectivity between the VWFA and language regions predicted reading, functional connectivity between the same regions did not (Chen et al., 2019). Considering past evidence and our current findings, we suggest that the patterns of VWFA functional connectivity may be an intrinsic, stable property of the brain. However future studies employing causal methodologies (e.g., interventions and longitudinal designs) will be needed to test this hypothesis.

It has been suggested that brain connectivity serves as a blueprint for the organization of category-selective regions (Hannagan et al., 2015). This idea was put forward by observations that category-selective regions in the VTC show specific connections with distant brain regions that are involved in the processing of the same category (Stevens et al., 2015). Longitudinal studies revealed that structural connectivity patterns were observed before the emergence of category selectivity itself (Saygin et al., 2012, 2016). Recent studies have discovered that specific functional connectivity patterns of category-selective regions are already in place in neonates (Kamps et al., 2020; Li et al., 2020). Specifically, (Li et al., 2020) found that in infants the general anatomical location of the VWFA (on a template brain) is functionally coupled with the language network. These findings support the view that this cortical region is prewired for language, and that the connectivity of this region determines the future emergence of the VWFA while children learn to read. Our findings lend further support to this notion by showing that subregions of the VWFA have distinct functional connections with distant regions that correspond to their respective role in word processing. Further, our findings reveal a striking overlap between the connectivity of frontal language regions and the boundary between VWFA-1 and VWFA-2. We suggest that the intrinsic link between frontal language regions and the occipitotemporal cortex may govern the emergence of VWFA-2. Future studies can examine this hypothesis directly by testing whether functional connectivity can predict the boundary between the more lexical VWFA-2 and the more visual VWFA-1 in larger samples.

The current study capitalizes on existing large-scale datasets, which come with their own advantages and limitations: NSD provides high-quality data, long scan durations and individually-defined ROIs, but has a small number of participants and no reading assessments. On the other hand, HBN provides a large developmental sample with reading assessments, but no individual text-selective ROIs. Addressing our question using these two complementary datasets allows us to leverage their strengths to provide a broad perspective on functional connectivity, as well as affirm the replicability of our findings. Future longitudinal studies are needed to delineate the concurrent developmental trajectory of category-selectivity, functional connectivity and structural connectivity.

## Acknowledgements

This work was supported by NICHD R01-HD095861 to JDY, Stanford Maternal and Child Health Research Institute award to IK, the National Science Foundation Graduate Research Fellowship (DGE-1656518) to EK and the Zuckerman-CHE STEM Leadership Program to MY.

