## Supplementary Figures1-7 for "The transition from vision to language: distinct patterns of functional connectivity for sub-regions of the visual word form area"

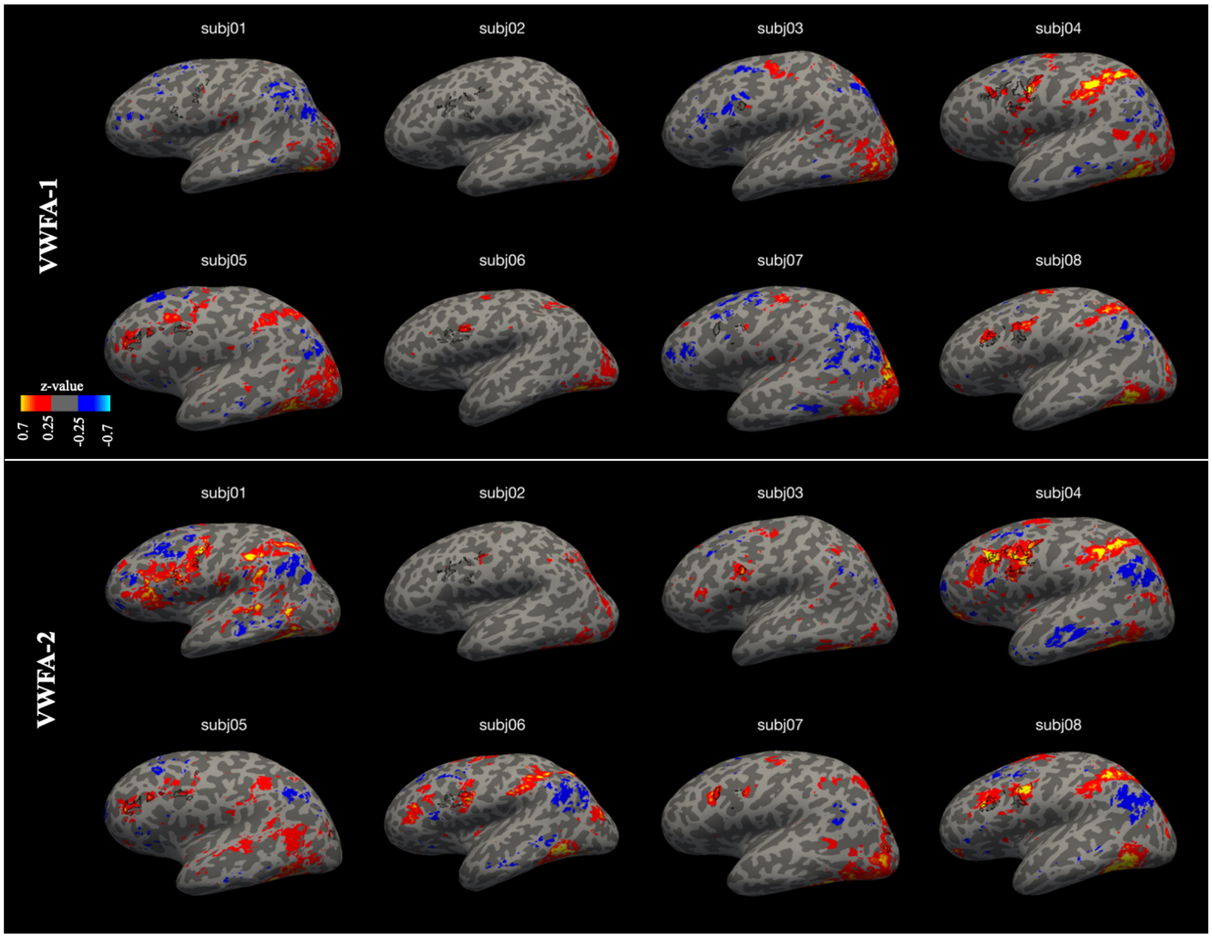


**Supplementary Figure 1:** Seed-based connectivity maps for all NSD subjects, left hemisphere (expands on Figure 2 from the main text)

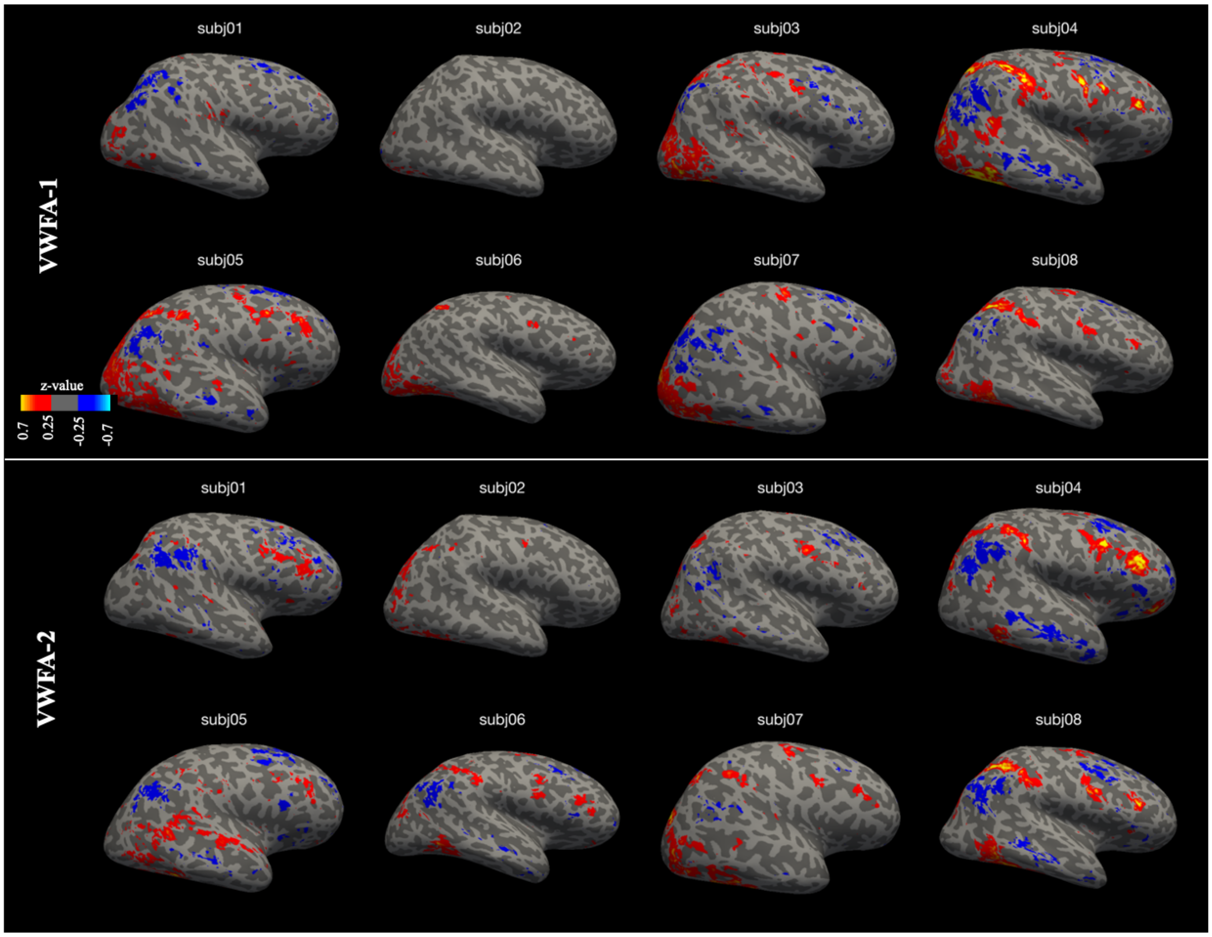


**Supplementary Figure 2:** Seed-based connectivity maps for all NSD subjects, right hemisphere (expands on Figure 2 from the main text)

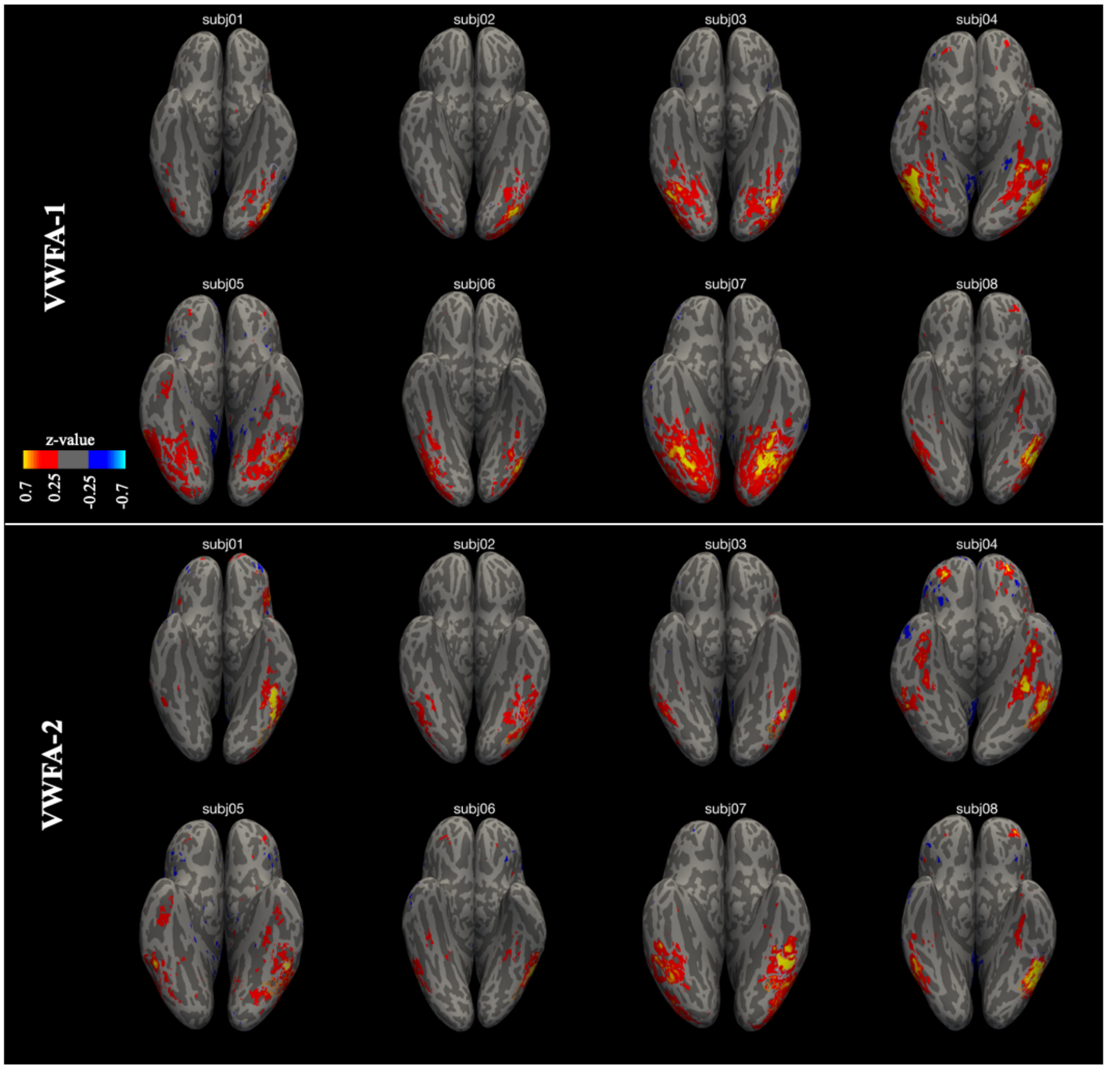


**Supplementary Figure 3:** Seed-based connectivity maps for all NSD subjects, inferior view (expands on Figure 2 from the main text)

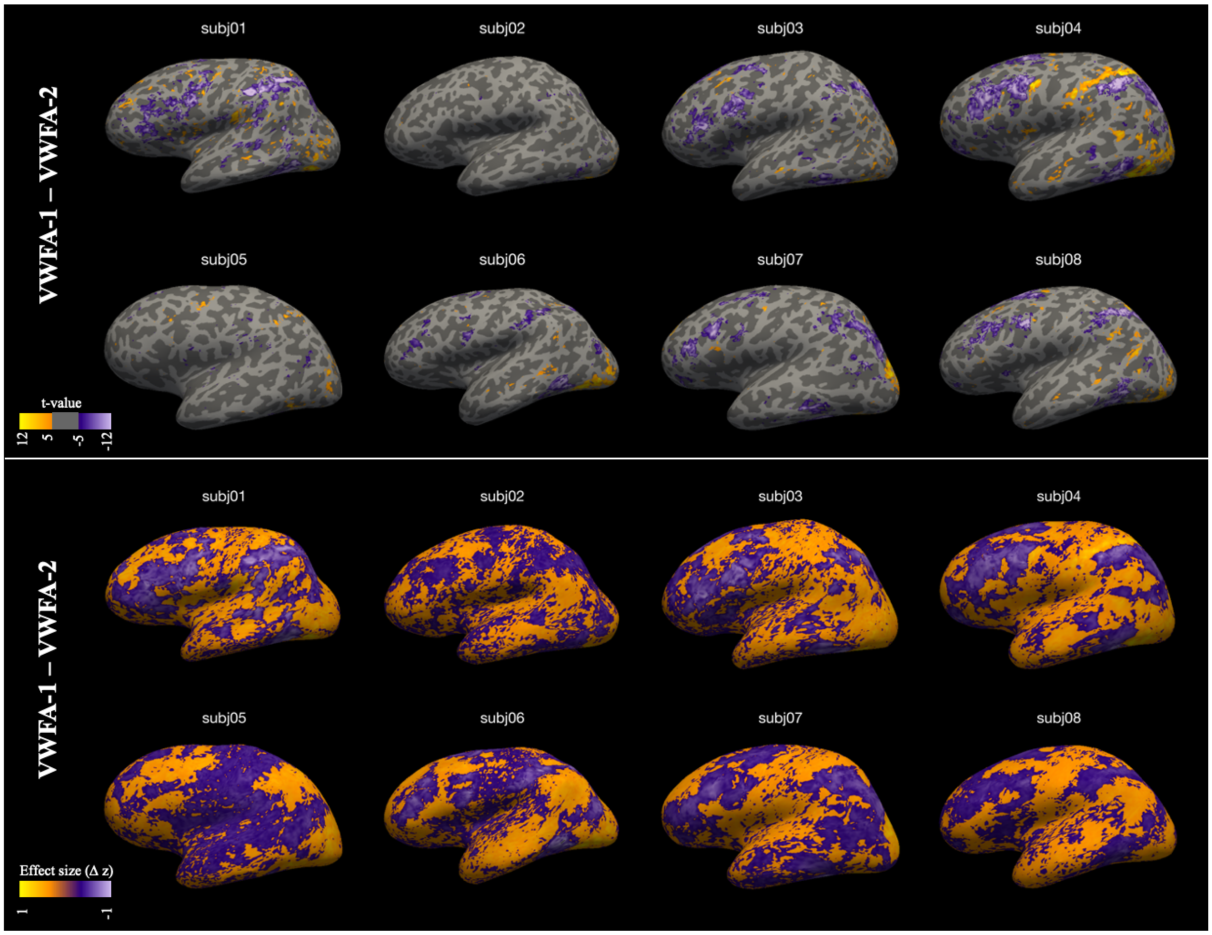


**Supplementary Figure 4:** Maps comparing the connectivity strength of VWFA-1 and VWFA-2 throughout the brain, for all NSD subjects. Top panel shows thresholded t-test maps, while the bottom panel shows the unthresholded effect size (z(VWFA-1) - z(VWFA-2). Left hemisphere

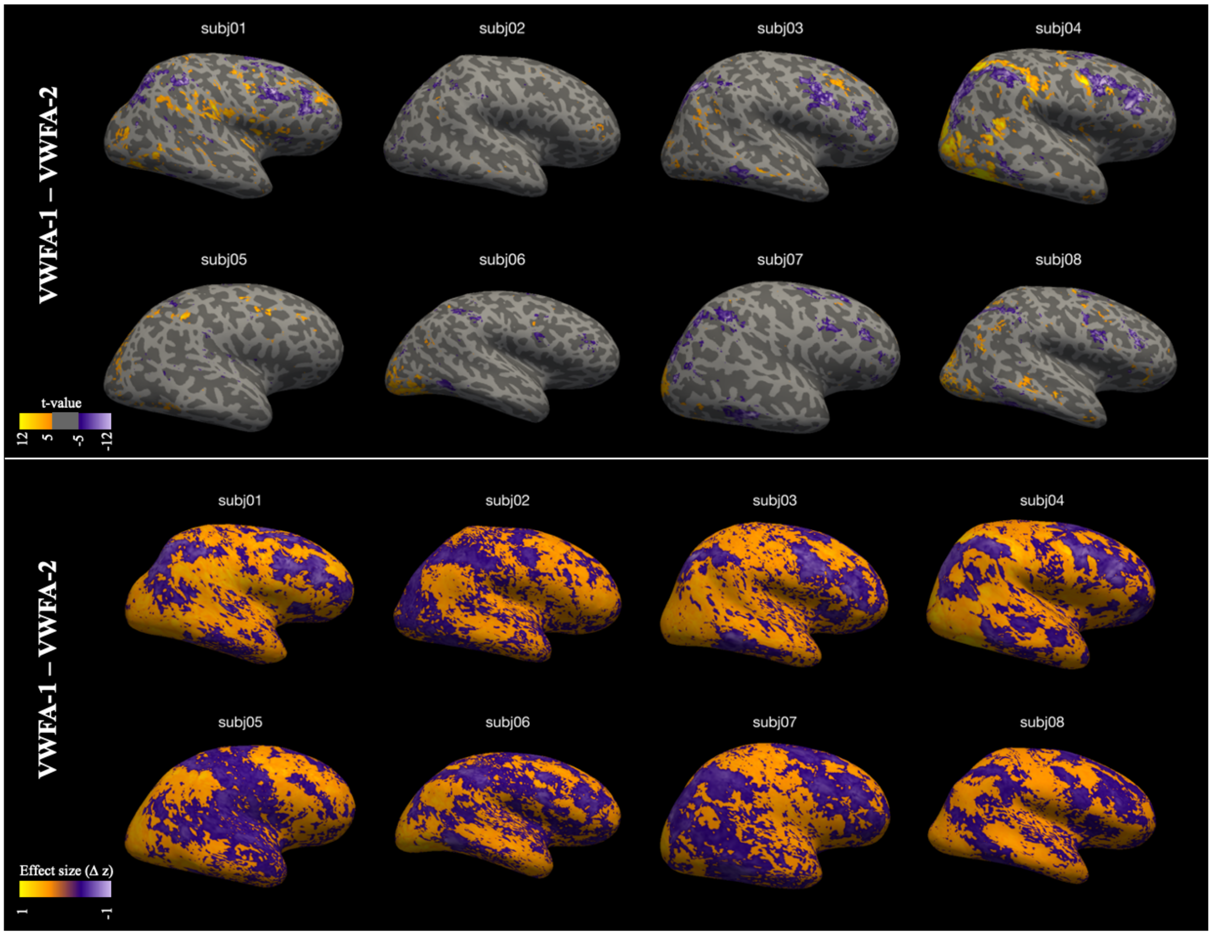


**Supplementary Figure 5:** Maps comparing the connectivity strength of VWFA-1 and VWFA-2 throughout the brain, for all NSD subjects. Top panel shows thresholded t-test maps, while the bottom panel shows the unthresholded effect size (z(VWFA-1) - z(VWFA-2). Right hemisphere

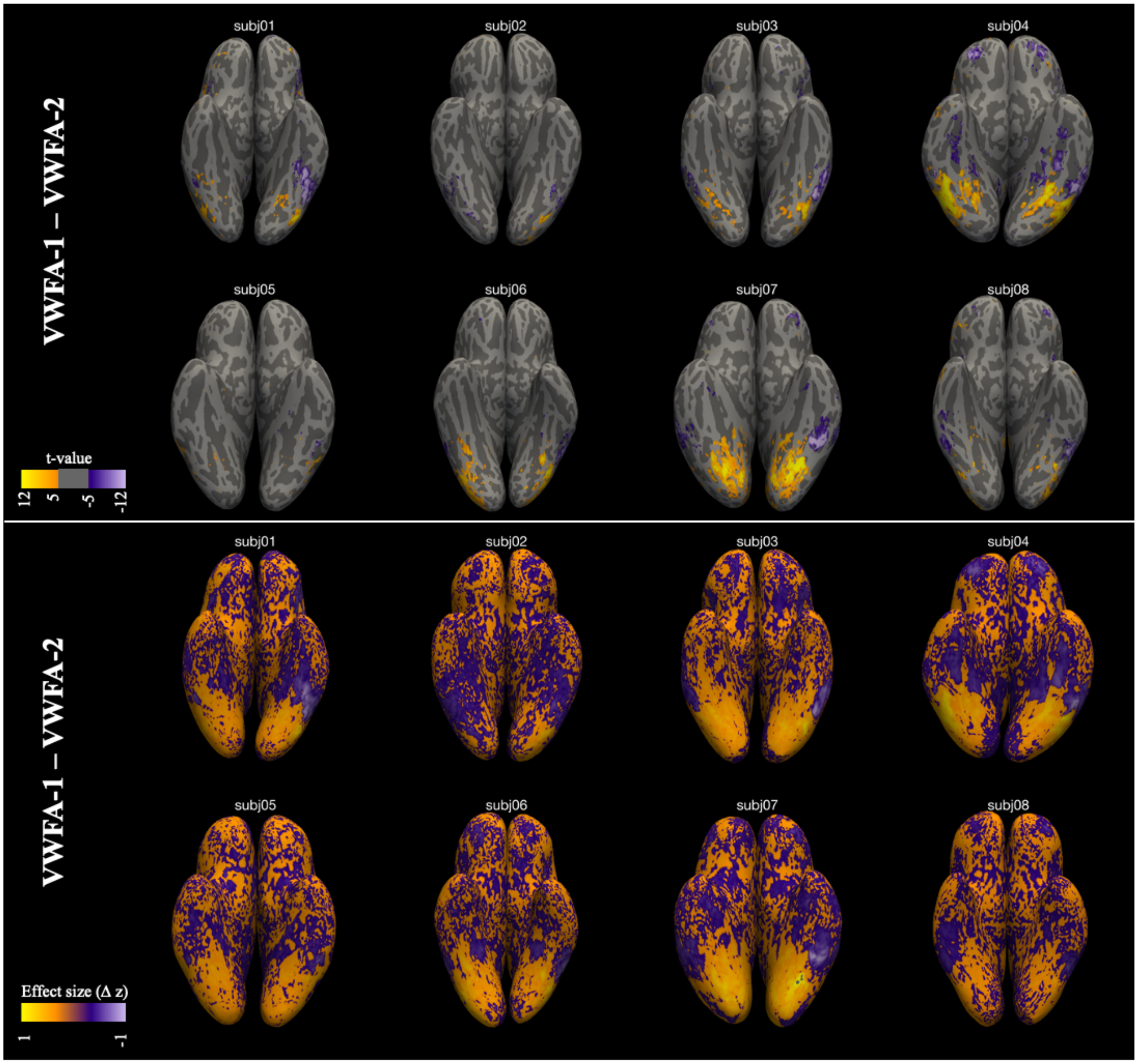


**Supplementary Figure 4:** Maps comparing the connectivity strength of VWFA-1 and VWFA-2 throughout the brain, for all NSD subjects. Top panel shows thresholded t-test maps, while the bottom panel shows the unthresholded effect size (z(VWFA-1) - z(VWFA-2). Inferior view

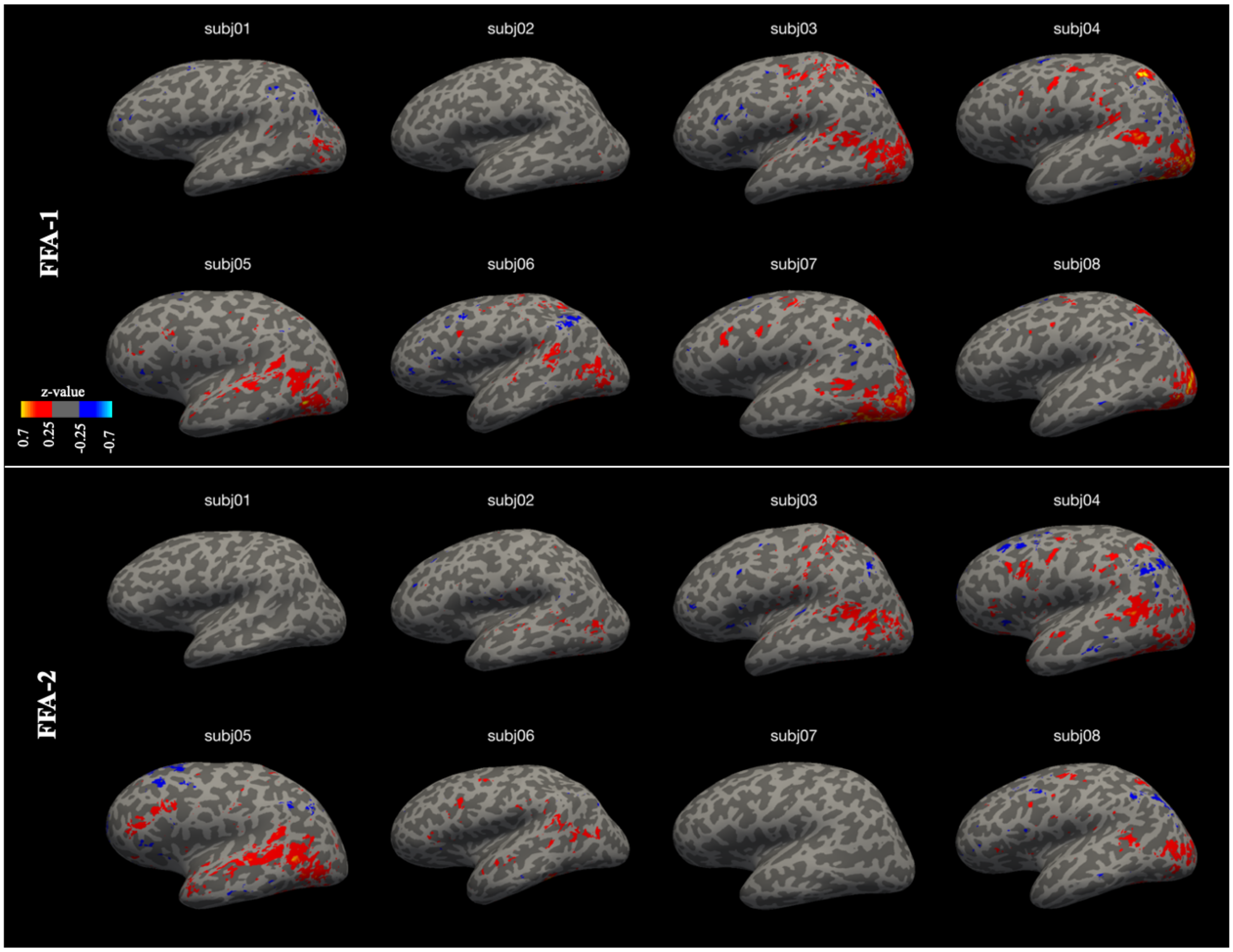


**Supplementary Figure 7:** Seed-based connectivity maps for face-selective ROIs (FFA-1/FFA-2) for all NSD subjects, left hemisphere (expands on Figure 7 from the main text)
